# NMR Mapping of Disordered Segments from a Viral Scaffolding Protein Encapsulated in a 23 MDa Procapsid Complex

**DOI:** 10.1101/539965

**Authors:** Richard D. Whitehead, Carolyn M. Teschke, Andrei T. Alexandrescu

## Abstract

Scaffolding proteins are requisite for the capsid shell assembly of many tailed dsDNA bacteriophages, some archaeal viruses, herpesviruses, and adenoviruses. Despite their importance, no high-resolution structural information is available for scaffolding proteins within capsids. Here we use the inherent size limit of NMR to identify mobile segments of the phage P22 scaffolding protein in solution and when incorporated into a ~23 MDa procapsid complex. Free scaffolding protein gives NMR signals from both the N and C-terminus. When scaffolding protein is incorporated into P22 procapsids, NMR signals from the C-terminal helix-turn-helix (HTH) domain disappear due to binding to the procapsid interior. Signals from the N-terminal domain persist, indicating this segment retains flexibility when bound to procapsids. The unstructured character of the N-terminus coupled with its high content of negative charges, is likely important for the dissociation and release of scaffolding protein, during the genome packaging step accompanying phage maturation.

**Graphical Abstract:** 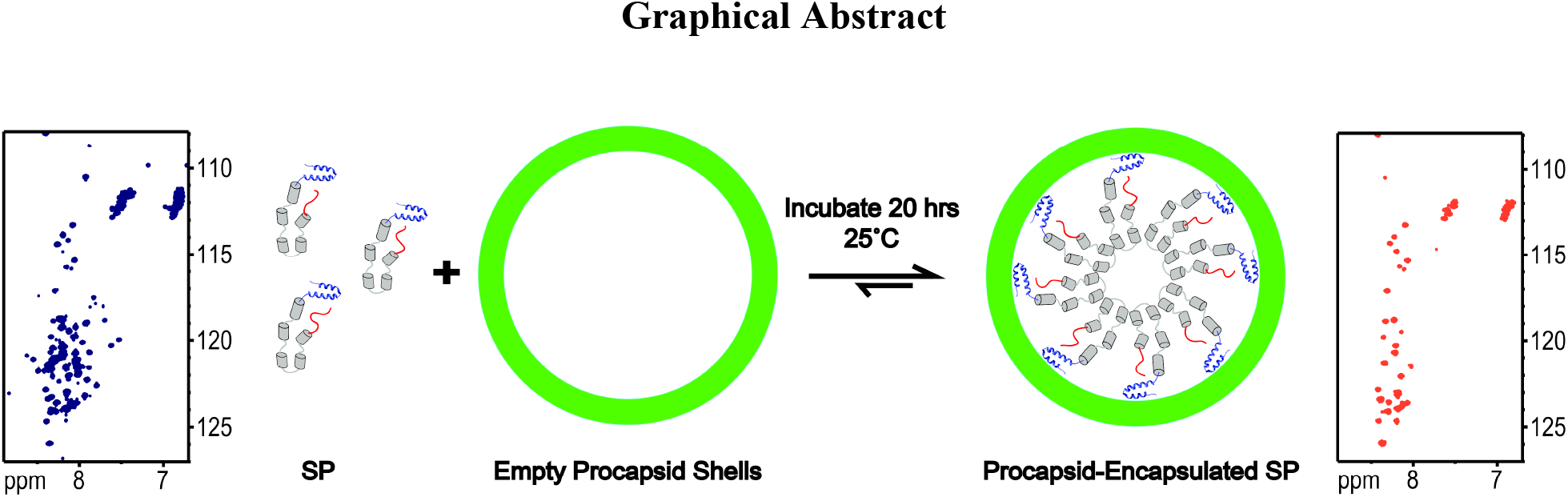

Scaffolding protein (SP) nucleates the assembly of phage P22 coat proteins into an icosahedral capsid structure that envelops the viral genome. NMR spectra of free SP show signals from the N-terminus (red) and a helix-turn-helix domain at the C-terminus (blue). When SP is incorporated into empty phage P22 procapsids to form a 23 MDa complex, the subset of signals from the N-terminal 40 residues persist indicating this segment is disordered. The unfolded nature of the N-terminus coupled with its negatively charged character, is important for the functional requirement of SP to exit the capsid as it becomes packaged with its genome.

## Introduction

In many viruses, scaffolding proteins (SPs) are required to ensure the correct organization of coat proteins (CPs) and other minor capsid proteins into a precursor structure, called a procapsid (Dokland, 1999; Fane and Prevelige, 2003). Even though SPs are critical for viral assembly, and therefore potential therapeutic targets (Parker et al., 1997a), their structural properties are poorly understood. Here we investigate the structure of full-length scaffolding protein from bacteriophage P22. Phage P22 is a well-characterized model for the class of tailed double-stranded DNA (dsDNA) bacterial viruses. Between 60-300 copies of a 303 amino acid SP (33.6 kDa) catalyze the assembly of 415 copies of CP (46.8 kDa) to form the icosahedral T=7 procapsid of phage P22 (~23 MDa) (Keifer et al., 2014a). The SPs are also required for incorporation of the dodecameric portal protein complex at one vertex of the icosahedron, as well as other minor internal proteins (Bazinet and King, 1988; Israel, 1977). SPs are not found in mature virions, as they are released without proteolysis during genome packaging. The SP is then recycled for subsequent use in multiple rounds of phage assembly (King and Casjens, 1974). Without SP, phage P22 coat protein fails to assemble into normal T=7 capsids. Instead, aberrant ‘petite’ T=4 procapsids or spiral structures are formed, both of which are non-infectious (Earnshaw and King, 1978; Thuman-Commike et al., 1998). Lack of proper assembly in the absence of SP is a general feature of this class of viruses.

Although the functional domains of SP are well-characterized (Greene and King, 1996, 1999; Weigele et al., 2005), its structural properties have proven elusive because of the unusual dynamic properties of the protein (Tuma et al., 1996). The amino acid sequence of SP has a high content of charged residues (31%) and a low content of hydrophobic residues, typical of intrinsically disordered proteins (IDPs) (van der Lee et al., 2014). There is a marked asymmetry in the distribution of charges in the SP amino acid sequence, giving rise to a highly acidic N-terminus, and basic C-terminus (Supporting Information Figure S1). This charge distribution suggests that SP belongs to the electrostatic hairpin group of IDP structures (van der Lee et al., 2014). Biochemical data are consistent with the sequence properties of SP. The protein exhibits a non-cooperative unfolding transition characteristic of molten globule conformations that have secondary structure coupled with a liquid-like fluctuating tertiary structure. SP also exhibits rapid amide proton hydrogen exchange typical of marginally stable secondary structure (Greene and King, 1999; Tuma et al., 1996). These properties are not unique for the SP of P22; conformational flexibility is a hallmark of viral scaffolding proteins, that appears to be required for their functionality (Dokland, 1999; Fane and Prevelige, 2003; Medina et al., 2011). Nevertheless, the SP of P22, as well as those from phages lambda, φ29, SPP1, and T4 have a high content of predicted and/or measured α-helical secondary structure (Lee and Guo, 1995; Mesyanzhinov et al., 1990; Parker et al., 1997a; SL et al., 2008). Moreover in the SP of phage P22, the C-terminal domain that is critical for CP binding during procapsid assembly adopts an independently folded helix-turn-helix (HTH) domain structure, as determined by NMR (Sun et al., 2000). Thus, while SP lacks a fixed global hydrophobic core, it is organized into distinct functional domains that are used to fulfill the protein’s multiple roles in the procapsid assembly process (Weigele et al., 2005). In previous biochemical cross-linking studies we saw evidence that P22 SP undergoes conformational changes during encapsulation into procapsids (Padilla-Meier and Teschke, 2011). Here, we use the intrinsic size limit of NMR to define the regions of SP that remain unfolded after assembly.

## Results

Figure 1 shows data from a pH titration of the phage P22 SP monitored by ^1^H-^15^N HSQC NMR spectra. Figure 2 shows the change in secondary structure of SP as a function of pH followed by far-UV circular dichroism (CD). At pH 2 the CD spectrum is characteristic of an unfolded random coil conformation (Johnson, 1988) (Figure 2) and ~300 ^1^H-^15^N correlations are seen in the NMR spectrum, consistent with the 303 amino acids in the sequence of SP (Figure 1). With increasing pH, the CD spectra increasingly show a profile consistent with α-helical structure as the SP protein folds (Figure 2). Concomitantly, crosspeaks broaden out in the ^1^H-^15^N HSQC NMR spectra and only about 100 peaks are left near neutral pH (Figure 1). Similar results were obtained with the related SP of phage CUS-3 (Figure S2 and S3).

**Figure 1.**
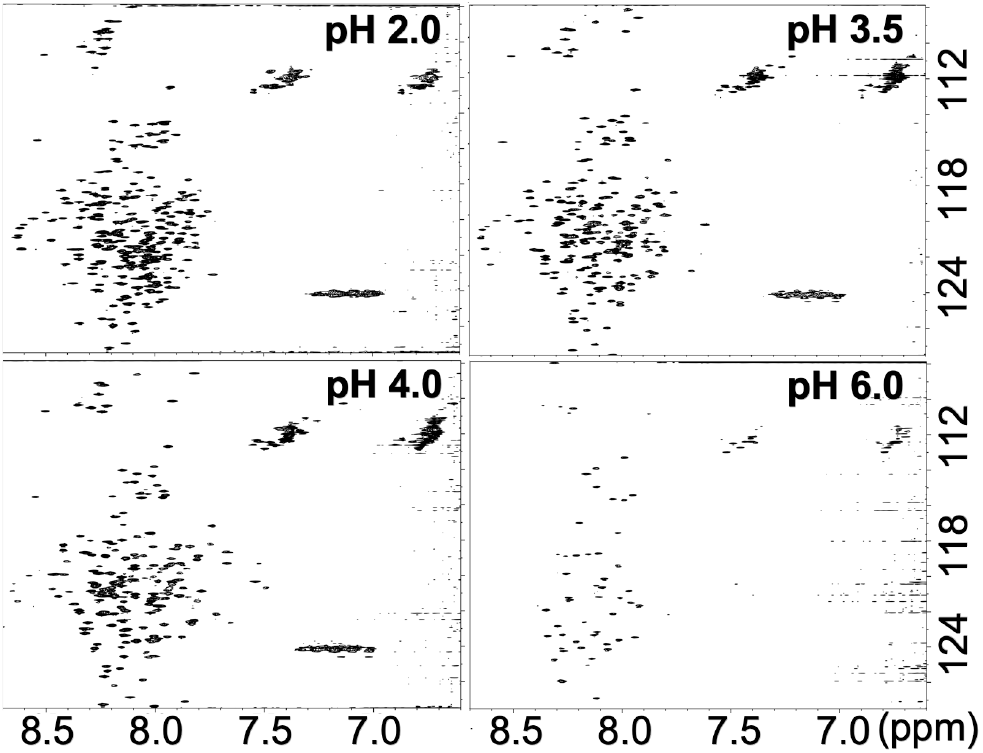
Representative ^1^H-^15^N HSQC spectra from a pH titration of P22 SP (115 μM) at a temperature of 30 °C.

**Figure 2.**
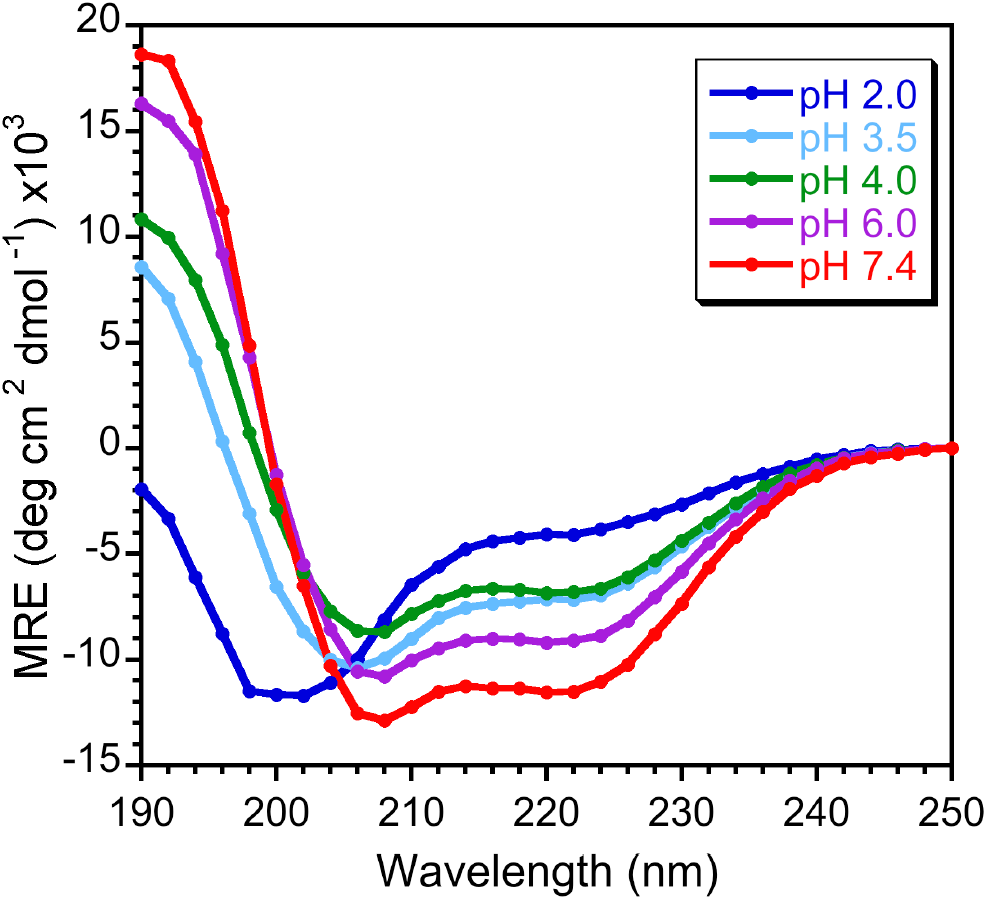
Far-UV CD of the SP from P22 (3 μM) as a function of pH. The data were obtained at 30 °C.

Sequence-specific NMR assignments for ^13^C/^15^N-labeled SP from phage P22 were obtained at pH 6.0 and a temperature of 35 °C using 3D NMR experiments (Figure S4 and Table S1). The ^1^H-^15^N correlations that survive at pH 6.0 (Figure 1) are mostly from the N-terminal residues 1-40 that have chemical shifts characteristic of an unfolded random coil conformation, and the C-terminal residues 264-303 that correspond to the folded HTH CP-binding domain of SP (Sun et al., 2000). Many of the ^1^H^N^ NMR signals from the C-terminal segment are shifted upfield in the NMR spectrum, consistent with the α-helical HTH NMR structure of this domain (Sun et al., 2000).

The majority of signals from the central portion between residues 41-263 of free SP were not detected in NMR spectra. Biophysical studies indicate that SP consists of loosely organized domains of alpha helical structure, connected by turn and random coil segments (Greene and King, 1999; Tuma et al., 1998; Tuma et al., 1996). Furthermore, SP exists in a rapidly reversible monomer-dimer-tetramer dynamic equilibrium where the dimers are thought to be the catalytically active species in procapsid nucleation (Parker et al., 1997b). Thus the broadening of NMR resonances from the 41-263 segment of SP (Figure 1) accompanying the folding transition to an α-helix structure at neutral pH demonstrated by CD (Figure 2), could be due to a combination of the following mechanisms. (1) NMR signals could be broadened beyond detection upon folding due to the large size of the protein (33.6 kDa for the monomer) and of its oligomeric complexes. (2) Exchange broadening could occur because of rapid interconversion between the monomer, dimer (K_d_ = 91 μM)(Parker et al., 1997b), and tetramer oligomerization states. (3) Exchange broadening could reflect a ‘molten globule’-like conformation for the 41-263 segment of SP, with α-helical secondary structure but no uniquely defined tertiary structure (Tuma et al., 1998; Tuma et al., 1996). We note that the NMR structure of the globular α-helical HTH domain of SP was determined at pH 4 (Sun et al., 2000). The CD data in Figure 2 indicate a near doubling of the ellipticity at 220 nm, characteristic of α-helical structure, between pH 4.0 and pH 7.4. It is therefore extremely unlikely that the transition to α-helical structure with increasing pH is entirely due to the HTH domain. Rather, a substantial amount of α-helical structure must be formed in the central 41-263 domain as SP becomes increasingly folded towards neutral pH.

We next used NMR to investigate the properties of SP when it is encapsulated into phage P22 procapsids (Figure 3). These experiments are conceptually analogous to “in-cell NMR experiments” (Croke et al., 2008; McNulty et al., 2006; Serber et al., 2006; Theillet et al., 2014), where signals from small proteins, or flexible segments of proteins, can be observed when they are incorporated inside living cells, as long as the isotope-labeled proteins of interest do not interact strongly with other large cellular components (Croke et al., 2008; McNulty et al., 2006; Serber et al., 2006; Theillet et al., 2014). We believe this “in-virus” NMR strategy could be more generally applicable to the study of the dynamic properties of macromolecules encapsidated into virus particles, including cargo molecules encased in viral capsids for nanotechnology applications. Additionally, such studies could assess the level of interaction of cargo molecules with the virus and probe the release properties of cargo.

**Figure 3.**
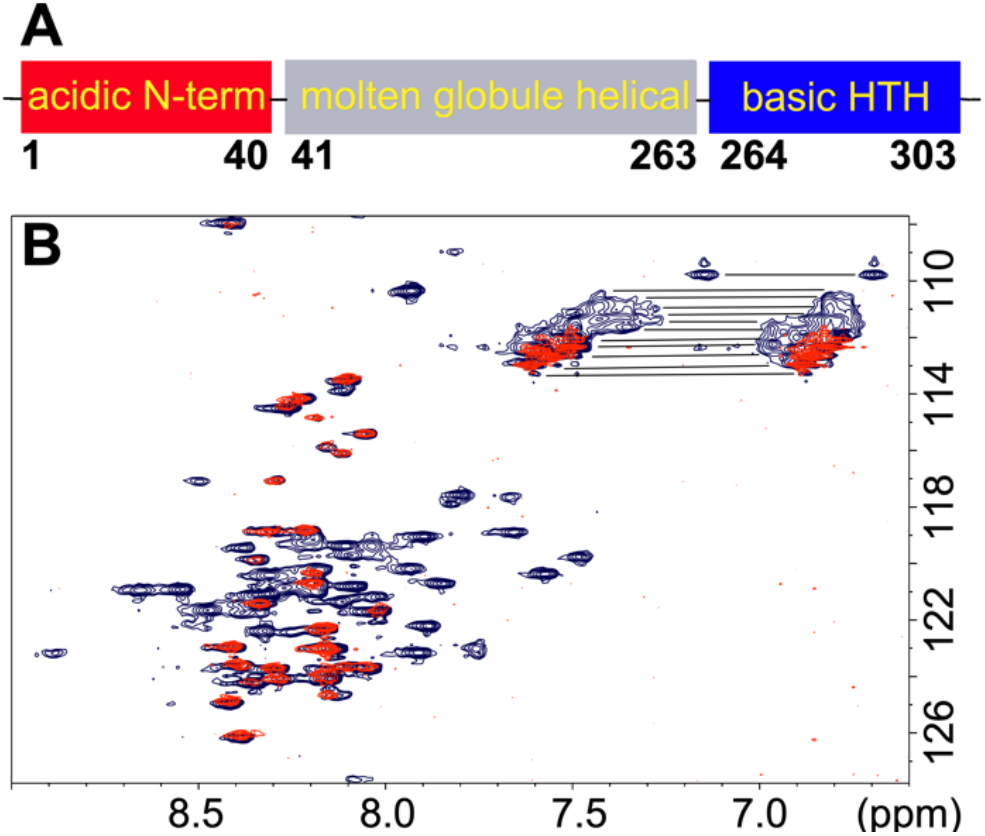
NMR of SP incorporated into phage P22 procapsids. (A) Domain diagram of the phage P22 SP protein. (B) Superposition of the ^1^H-^15^N HSQC spectra of free SP (blue), with SP in the 23 MDa procapsid complex (red). For NMR assignments see Figure S5 and S6. NMR data were acquired on samples in 20 mM sodium phosphate, 50 mM NaCl, pH 7.0, at a temperature of 35 °C. Horizontal lines in the upper right corner indicate unassigned crosspeaks from asparagine and glutamine side-chain NH_2_ groups.

^15^N-labeled SP was incorporated into unlabeled empty procapsid shells as previously described (Fuller and King, 1981; Prevelige et al., 1988). SP was mixed with procapsid shells at a molar ratio of 60 SP to 1 empty procapsid shell, to allow rebinding of SP to the interior of the procapsids (Greene and King, 1994; Teschke et al., 1993). The 60:1 ratio of SP:procapsid was chosen to ensure tight binding of SP to the procapsid. Higher ratios of up to 300:1, result in a mixture of strongly and weakly bound species (Parker et al., 2001). The encapsulated SP was separated from any unbound protein by ultracentrifugation. This yielded a capsid-SP complex of between 22-24MDa, depending on the amount of incorporated scaffolding protein (Keifer et al., 2014b). Sedimented procapsids with encapsulated ^15^N-SP were resuspended in phosphate buffer and NMR data were collected. All of the ^1^H-^15^N HSQC crosspeaks from the procapsid-encapsulated SP (shown in red in Figure 3B) are from residues 4-40 in the acidic N-terminal domain of SP (shown in red in Figure 3A). In the spectrum of free SP under identical conditions (blue contours in Figure 3B), most of the assigned correlations (Table S1, Figure S5 and S6) are from the disordered N-terminal domain (residues 4-40) also seen in the encaplsulated SP, and the basic folded HTH domain (residues 264-303) shown in blue in Figure 3A. NMR signals from the HTH domain broaden beyond detection in the encapsulated SP due to its tight binding to the capsid. The central portion of SP (residues 41-263) correpsonding to the α-helical oligomerization domain (gray in Figure 3A) shows no crosspeaks in the procapsid-encapsulated SP, and only 10-20 crosspeaks in the free protein. The lack of NMR signals for the large central portion of SP in the free protein is likely due to its molten globule character, which is often associated with conformational exchange broadening of protein NMR signals (Alexandrescu et al., 1994; Alexandrescu et al., 1993; Redfield, 2004). Similar results to those presented in Figure 3 were obtained with a variant of SP that contained an N-terminal His6-tag, to allow higher protein purification yield (Figure S7 and S8). Control experiments in which 15N-labeled SP was pelleted with the procapsids were done to verify that the observed NMR signals were from encapsulated SP rather than the free protein (Figure S9 and S10).

## Discussion

During maturation of the procapsid to a virion, there are large structural rearrangements of the N-arm and A-domain of CP, leading to rearrangements of the internal procapsid surface that results in a switch from a predominantly negative to predominantly positive character. This switch in the electrostatic properties of the internal capsid surface, likely triggers exit of SP, as Coulombic interactions between SP and CP present in the procapsid are broken upon maturation (Kang and Prevelige, 2005; Parent et al., 2010; Teschke and Parent, 2010). Previous studies of SP showed that the N-terminal domain is important for the exit of SP from procapsids, and that removal of the N-terminus yields virions with an incomplete dsDNA genome (Weigele et al., 2005). We hypothesize the highly negative N-terminal domain could compete for interactions between the negatively charged surface of CP and the positively charged HTH domain, thereby facilitating exit of SP.

In summary, we observe NMR signals from the first 40 amino acids of phage P22 SP when it is encapsulated into phage P22 procapsids. Signals from the basic C-terminal domain of SP disappear, because it is tightly bound to CP assembled to form a procapsid. That NMR signals from the N-terminal domain persist indicates that this segment is unfolded in the SP-procapsid complex. The intrinsic disorder of the N-terminal segment is likely to have biological significance, as this segment has been shown to be necessary for SP exit during viral maturation. The present results show that the well-known size limitation of NMR can be used to advantage, as a filter to identify disordered segments even in very large supramolecular complexes of proteins. In this way, NMR can provide a unique perspective on dynamic and disordered elements of macromolecules not accessible by other techniques.

## Star*Methods

Detailed methods are provided in the online version of this paper and include the following:

- Protein Expression and Purification
- Encapsulation of SP into Procapsids
- NMR Spectroscopy
- Circular Dichroism Spectroscopy

## Supplemental Information

Supplemental Information includes five figures and one video and can be found with this article online at: ***

## Author Contributions

C.M.T and A.T.A conceived the study. R.D.W. purified proteins, preformed CD and NMR experiments, and analyzed data with A.T.A. A.T.A., R.D.W. and C.M.T. wrote the manuscript.

## Experimental Procedures

### Expression and purification of proteins for NMR

The wild type P22 scaffolding protein (Pedulla et al., 2003) (lacking a His_6_ tag), henceforth ‘SP’, was prepared according to the following procedure. The SP containing pET plasmid was transformed into BL21 (DE3) cells for protein expression. Cells were grown in M9 minimal medium supplemented with 1 g/l ^15^NH_4_Cl to make ^15^N-isotope enriched protein samples. Following inoculation of the M9 media with a starter culture grown overnight, the cells were grown at 37°C to an OD_600_ of 0.5, at which point SP expression was induced with 1 mM IPTG (Gold Biotechnology, St Louis, MO). The cells were grown for a further 16 h at 18 °C. Cells were harvested by centrifugation at 7,800 g for 10 minutes in an F9 6×1000 LEX rotor (Thermo Scientific, Waltham MA), using a Lynx 6000 centrifuge (Thermo Scientific, Waltham MA), The cell pellet was resuspended in 20 mM sodium phosphate buffer. Cells were lysed using a Misonix sonicator (Misonix, Farmingdale NY), operating at an amplitude of 37, with 15 s pulses between 30 s intervals, for a total time of 3 min. Cell debris was sedimented by centrifugation for 15 min using an F18-12×50 rotor (Thermo Scientific, Waltham MA) operating at 31,920 g in a Sorvall RC6+ centrifuge (Thermo Scientific, Waltham MA). The supernatant was placed in a 65°C water bath for 10 minutes, as a purification step to heat-denature the bulk of *E.coli* proteins other than SP, which is able to refold after high heat treatment (Greene and King, 1999). The solution was then chilled on ice for 5 min, and spun at 31,920 g for 10 minutes in a Sorvall RC 6+ (Thermo Scientific, Waltham MA) centrifuge using an F18-12×50 rotor (Thermo Scientific, Waltham MA) to pellet the heat-precipitated proteins. The supernatant was loaded on an SP-Sepharose fast flow cation exchange column (GE Heathcare, Chicago Il), run at 1 ml /min using a 0-100% linear gradient from 0 M to 1 M NaCl, in 20 mM sodium phosphate, pH 7.4. Fractions containing purified SP (monitored by SDS-PAGE) were concentrated using an 80% saturated solution of ammonium sulfate (516 g/l), and dialyzed against 20 mM sodium phosphate, pH 7.4. Protein samples were stored at −80°C.

The purification scheme described above for wild type SP was used to determine that a variant of SP with an N-terminal His_6_-tag followed by a thrombin protease cleavage site for protein purification (Cortines et al., 2011) (together the tag is ~20 a.a.), henceforth denoted His_6_-SP, does not interfere with packaging of SP into procapsids or NMR spectra. Note that cleavage of His6-SP by thrombin leads to non-specific proteolysis within the interior of the disordered SP protein, so that this approach to remove the tag could not be used. With the His6-SP construct, yields were on the order of 50-70 mg of pure protein per liter of cell culture, or about 2-3 times larger than the ~25 mg/l yield for wild type SP obtained by the purification scheme described above. Thus to obtain NMR assignments we used ^15^N/^13^C-labeled His6-SP since the yield with this construct was better than for wild-type SP. ^1^H-^15^N HSQC spectra of SP and His_6_-SP were virtually identical in both the free and capsid bound states (see Figs. S5-S8). The additional amide proton NMR signals expected from the extra amino acids in the N-terminal His_6_-tag and thrombin site of His_6_-SP were not seen in ^1^H-^15^N HSQC spectra. Presumably this is due to fast hydrogen exchange of amide protons from the tag segment. In our experience, NMR signals from amide protons in His_6_-tags are often missing in other proteins, regardless of whether they are folded or unfolded.

His_6_-SP was produced using a construct containing the P22 SP gene sub-cloned into the pET15b vector (Novagen, Madison, WI). The His_6_-SP in a PET vector was transformed into *E. coli* BL21(DE3) cells (New England Biolabs, Ipswich, MA) and expressed and purified as previously described for SP (REF). The exception was that after collection of cells by sedimentation, the cell pellet was resuspended in 20 mM HEPES, 300 mM NaCl, 10 mM imidazole, pH 7.4. The cells were lysed in the same manner as above. The supernatant was loaded onto a Ni-NTA agarose affinity column (Qiagen, Hilden, Germany), washed with ~30 mL of 20 mM HEPES, 300 mM NaCl, 10 mM imidazole, and eluted using a 0-80% linear gradient of a 20 mM HEPES, 300 mM NaCl, and 500 mM imidazole at pH 7.4. Fractions containing His_6_-SP were identified by SDS-PAGE.

### Encapsulation of SP into phage P22 procapsids

Empty procapsid shells were generated as previously described by treatment with 0.5 M GuHCl (Greene and King, 1994). ^15^N-labeled P22 SP (or His_6_-SP) was encapsulated into empty procapsid shells in a ratio of 60:1, to ensure tight binding of SP to the shell (Parker et al., 2001). Specifically, empty procapsid shells at a concentration of 0.86 μM were mixed with 52 μM SP, in 20 mM sodium phosphate containing 50 mM NaCl. The mixture was incubated for 20 h at 25 °C to allow incorporation of SP into the procapsids. To collect the complex, the sample was spun at 175,000 g for 20 min using a RP80AT rotor (Thermo Scientific, Waltham MA), in a Sorvall RC M120EX ultracentrifuge (Thermo Scientific, Waltham MA). The resulting pellet was then resuspended in 500 μl of 20 mM sodium phosphate (pH 7.6) containing 50mM NaCl, on a reciprocal shaker (Eberbach, Ann Arbor MI) operating at 180 osc/min, for ~20 hours at 4° C (Suhanovsky and Teschke, 2011). Samples were applied to 2.2 ml of 5-20% (w/w) linear sucrose gradients, to verify the presence of SP inside shells. Sucrose gradients were centrifuged at 104,813 g for 35 min at 20°C in a M12EX (Thermo Scientific, Waltham MA) centrifuge with a RP55S rotor (Thermo Scientific, Waltham MA). Gradients were fractionated into 100 μL aliquots, and the presence of SP inside shells was verified by 10% SDS-PAGE (Prevelige et al., 1988).

### NMR spectroscopy

NMR data were collected on a Varian Inova 600 MHz spectrometer equipped with a cryogenic probe (Agilent, Palo Alto, CA). To investigate the effects of pH, ^1^H-^15^N HSQC spectra were collected at a temperature of 30 °C on 115μM and 280 μM samples of P22 and CUS-3 His_6_-SP, respectively. For ^15^N-P22 SP encapsulated into procapsids, we found that a temperature of 35 °C at pH 7.0 gave optimal NMR spectra. The concentration of P22 ^15^N-SP (or His_6_-SP) for these spectra was ~300 μM, at a SP:capsid ratio of 60:1. The capsid-encapsulated ^15^N-SP samples were in 20 mM sodium phosphate buffer containing 50 mM NaCl. To obtain NMR assignments we used a 0.5 ^13^C/^15^N-labled sample of His_6_-SP, since the yield for the His_6_-tagged protein was better than for wild type SP. NMR assignments for His_6_-SP were readily translatable to those for wild type SP, since the ^1^H-^15^N HSQC spectra of the two variants were superposable. NMR data for assignments were recorded at a temperature of 35 ^ω^C, using a 0.5 mM protein sample in 20 mM sodium phosphate, pH 6.0. The 3D spectra used to obtain assignments included 3D HNCACB, HNCACO, HNCO, ^15^N TOCSY-HSQC, and ^15^N NOESY-HSQC (Cavanagh et al., 1996). NMR data were processed using the programs Felix 2001 (Felix-NMR, San Diego, CA) and NMRpipe (Delaglio et al., 1995). Spectra were analyzed with CCPNmr (Vranken et al., 2005) on the NMRbox platform (Maciejewski et al., 2017). DSS was used as an internal reference for ^1^H chemical shifts. ^13^C and ^15^N nuclei were referenced indirectly as described in the literature (Wishart et al., 1995).

### Circular Dichroism (CD) spectroscopy

CD spectra were recorded on an Applied Photophysics Pi-Star 180 spectropolarimeter (Surrey, UK) using a 1 mm path-length cuvette, at 30 °C. Final SP concentrations were 3 μM in 10 mM sodium phosphate buffer at varying pH values. Wavelength scans were collected between 190-250 nm using a 2 nm bandwidth, 2 nm step size, 30 s/point data averaging, for a total scan time of ~15 min.

