## Supplemental Information file for "NMR Mapping of Disordered Segments from a Viral Scaffolding Protein Encapsulated in a 23 MDa Procapsid Complex"

Richard D. Whitehead III[†], Carolyn M. Teschke[†‡*], and Andrei T. Alexandrescu[†*].

[†]Department of Molecular and Cell Biology and [‡]Department of Chemistry, University of Connecticut, Storrs Connecticut, 06269, United States.

### Supplemental Information

## P22 SP

```
MEPTTEIQAT EDLTLSGDHA AASADSLVVD NANDNAGQEE GFEIVLKDDE TAPKQDPAKN
AEFARRRIER KRQRELEQQM EAVKRGELPE SLRVNPDLPP QPDINAYLSE EGLAKYDYDN
SRALAAFNAA NTEWLMKAQD ARSNAVAEQG RKTQEFTQQS AQYVEAARKH YDAAEKLNIP
DYQEKEDAFM QLVPPAVGAD IMRLFPEKSA ALMYHLGANP EKARQLLAMD GQSALIELTR
LSERLTLKPR GKQISSAPPA DQPITGDVSA ANKDAIRKQM DAAASKGDVE TYRKLKAKLK
GIR
```

303 a.a.; 6% P (17/303); overall pI = 5.2 (28E+23D, 20R+22K)
Resid 1-130, pI = 4.4 (16E+12D, 9R+6K); Resid 131-303, pI = 9.2 (12E+11D, 11R+16K)
Resid 1-40, pI = 3.5 (5E+5D, 0R+0K); Resid 264-303, pI = 9.8 (1E+5D, 3R+7K)

### CUS-3 SP

```
MDQMAENTPE VEIETDASEQ IPDDVELAEK VETEDGSESS GNDAEEATET DDDESEQEFY
FGDEKLDSPT SEDGAEHGLV KHLRKTIKEK DRELKELMRQ SQKPVEQQPV ITQPPRMPKL
DDEDIGFDEE IYQQRMAKWA EDNGKYQQQE MARKQKEQEL QAAYQERLSK YQQRVKALKV
PGYQEAEQAV LEEIPIETQN AILFESEKPE IVVLALGRNA ELRKQLAEAT NPVAIGRLLE
RIESKARIMP KAKTTAATTP TVKGSNGAVI NNLDKLKAKA LETGDWTPYF AAKKAKK
```

297 a.a.; 5% P (15/297); overall pI = 4.7 (44E+20D, 13R+29K)
Resid 1-130, pI = 4.0 (26E+17D, 4R+9K); Resid 131-297, pI = 9.6 (18E+3D, 9R+20K)
Resid 1-40, pI = 3.4 (10E+5D, 0R+1K); Resid 260-297, pI = 9.9 (1E+2D, 0R+8K)

### SF6 Scaff

```
MENELIIDGQ VIDLSETQEN AEETIIQTES QPENESQDDN GKEVATEPEK TEETPEDYAL
RIGDEEIQLN ADDDDHIDGQ PAPQWVKDLR KGFKETQKEN RELRRQLEEA LAKPAEHQQP
QPDAIPPKPT LESCDYDEQA FEQALTDWHE KKGRVEQQQQ QKLRQQQEYQ QRFQQRVEAH
KQRAAKLPVK DYQEMEAIVL SELPPIQQEI IIHCADEGSE LLAYGLGKSQ QLRQRVAAET
DPIRAAFLLG QISKQVSLAP KPKKAIKPEP EVRGGGADAK QDEFNKLCPG AKIE
```

294 a.a.; 7% P (20/294); overall pI = 4.6 (41E+21D, 14R+23K)
Resid 1-130, pI = 4.2 (23E+13D, 5R+8K); Resid 131-294, pI = 6.0 (18E+8D, 9R+15K)
Resid 1-40, pI = 3.1 (9E+4D, 0R+0K); Resid 260-294, pI = 8.9 (4E+2D, 1R+7K)

### HSV-1 SP (UL26.5)

```
MNPVPTSGTP APAPPGDGSY LWIPASHYNQ LVAGHAAPQP QPHSAFGFPA AAGAVAYGPH
GAGLSQHYPP HVAHQYPGVL FSGPSPLEAQ IAALVGAIAA DRQAGGQPAA GDPGVRGSGK
RRRYEAGPSE SYCDQDEPDA DYPYYPGEAR GGPRGVDSRR AARQSPGTNE TITALMGAVT
SLQQELAHMR ARTSAPYGMY TPVAHYRPQV GEPEPTTTHP ALCPPEAVYR PPPHSAPYGP
PQGPASHAPT PPYAPAACPP GPPPPPCPST QTRAPLPTEP AFPPAATGSQ PEASNAEAGA
LVNASSAAHV DVDTARAADL FVSQMMGAR
```

329 a.a.; 18% P (58/329); overall pI = 6.1 (13E+11D, 17R+1K)
Resid 1-130, pI = 7.1 (3E+3D, 5R+1K); Resid 131-329, pI = 5.4 (10E+8D, 12R+0K)
Resid 1-40, pI = 5.9 (0E+1D, 0R+0K); Resid 292-329, pI = 4.4 (2E+3D, 2R+0K)

**Fig. S1 Sequences, proline residues fractions, and charge distributions of SPs.** Proline residues are colored green, acidic residues red, and basic residues blue. Theoretical p*I* values are shown for the proteins overall, and the indicated protein fragments (Casjens et al., 2004; Davison, 2011; King et al., 2007; Pedulla et al., 2003). The phages P22, CUS-3, and Sf6 are representative of three major groups from a family of over 150 P22-like phages (Casjens and Thuman-Commike, 2011). The sequence of the more distantly related SP from Herpesvirus-1 (Dokland, 1999) is included for comparison.

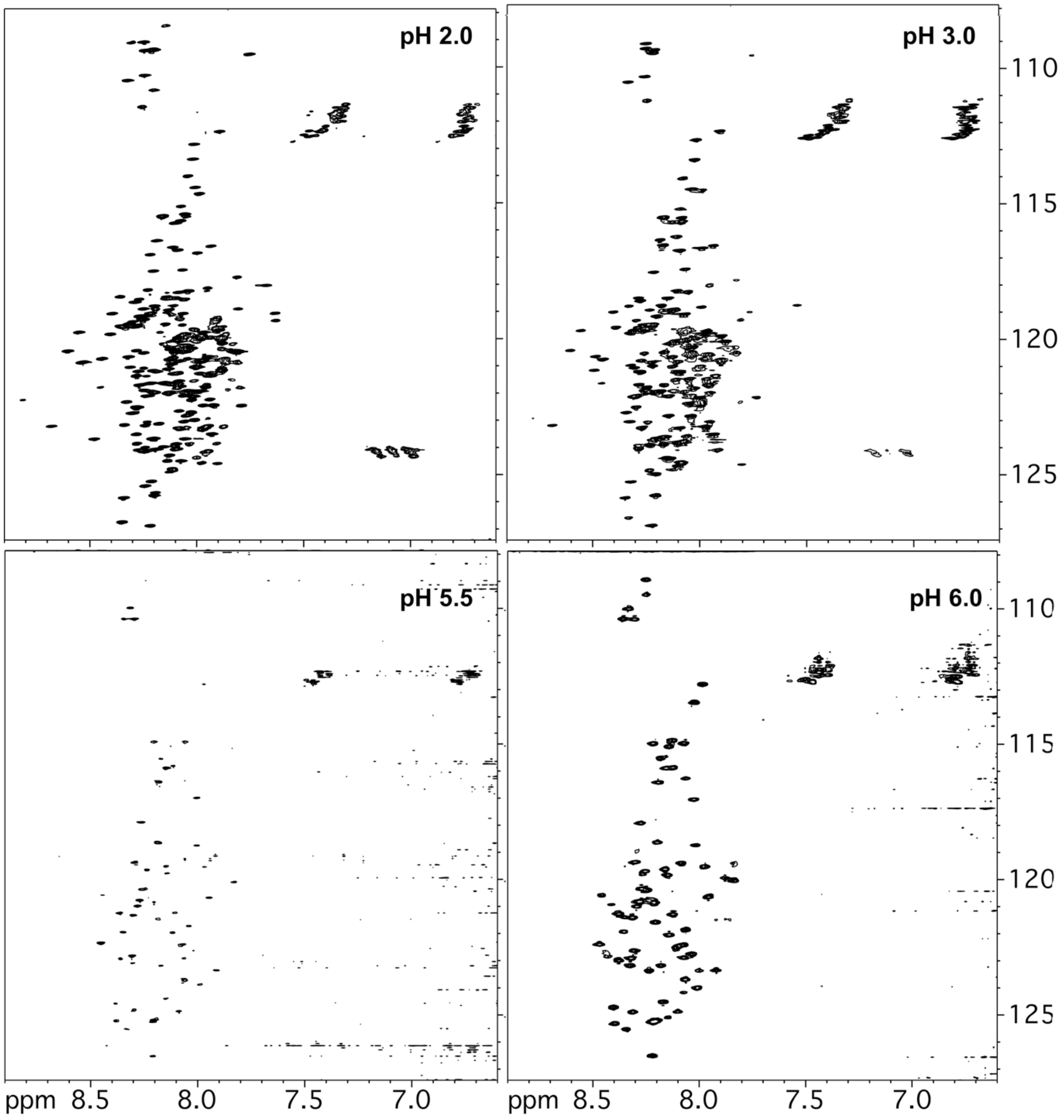

**Fig. S2 $^{1}$H-$^{15}$N HSQC spectra of CUS-3 His$_6$-SP as a function of pH.** The data were recorded on 280 μM protein samples at a temperature of 30 $^{o}$C. With increasing pH, crosspeaks are lost as residues that are initially unstructured at pH 2 become involved in the molten globule α-helical structure of the central domain of the protein. Thus while ~300 crosspeaks are seen at pH 2 when the protein is unfolded, only ~100 crosspeaks persist when the central domain adopts a molten globule α-helical structure near neutral pH. The enhanced broadening observed at pH 5.5 is due to increased aggregation, near the theoretical p*I* of 4.7 for the CUS-3 protein.

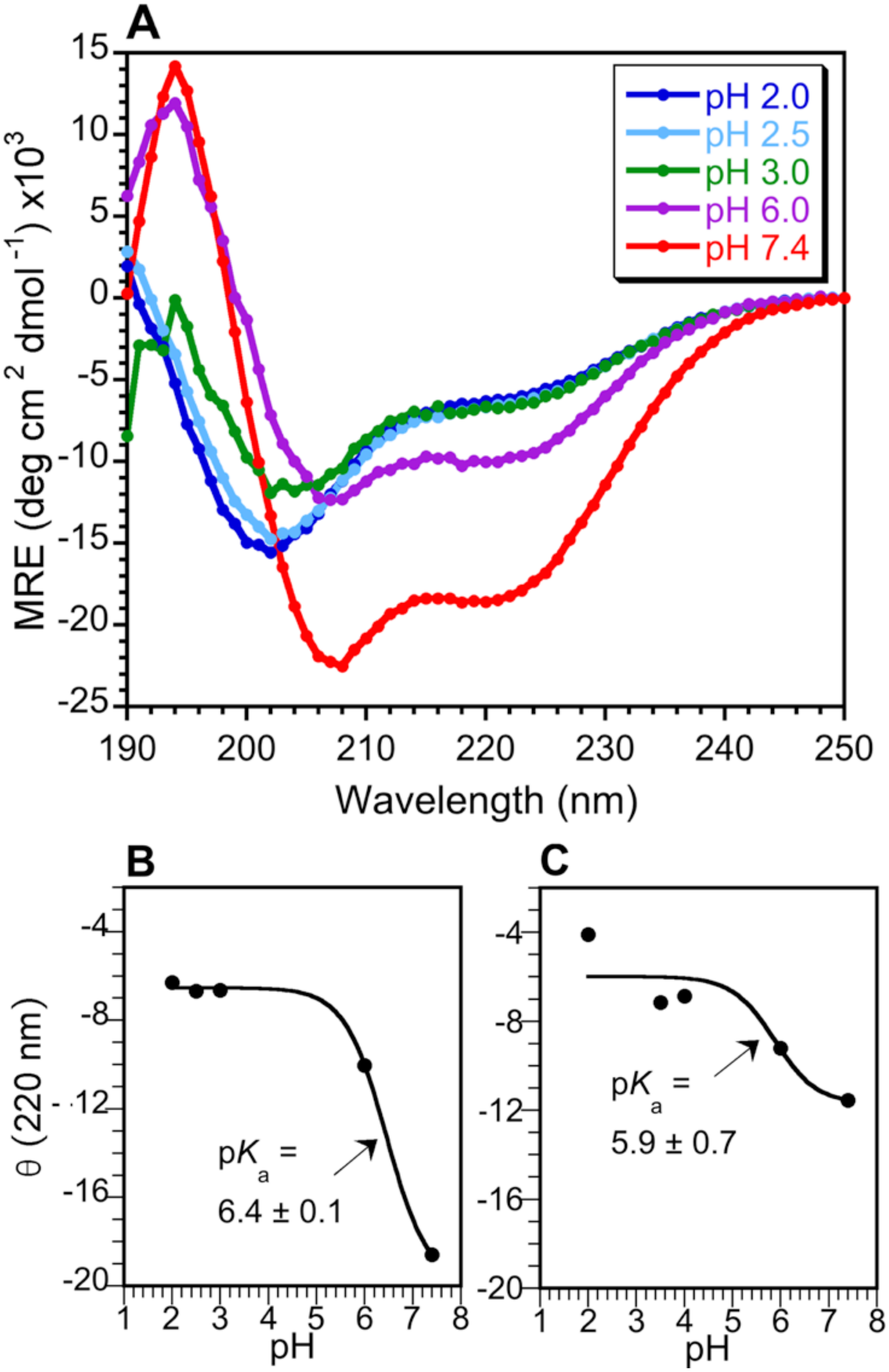

**Fig. S3 Increase of α-helix structure in His$_6$-SP with increasing pH. (A)** Far UV circular dichroism (CD) spectra of the CUS-3 His$_6$-SP as a function of pH. At pH 2, the spectrum is typical of a random coil. At pH 7.4 there is a large increase in the amount of α-helical structure as monitored by ellipticity at 208 and 220 nm. Data were collected on 3 µM samples of CUS-3 His$_6$-SP in 10 mM sodium phosphate buffer, 30 $^{o}$C. (**B**) Ellipticity at 220 nm for CUS-3 His$_6$-SP as a function of pH. Data from A, fit to an apparent p$K_a$ of 6.4. (**C**) P22 His$_6$-SP ellipticity at 220 nm as a function of pH. The data from Fig. 2 of the main text, fit to an apparent p$K_a$ of 5.9. The apparent p$K_a$ values obtained for both the P22 and CUS-3 His$_6$-SPs are close to the theoretical p*I* values for the central segments of the respective proteins: p*I* of 4.9 for CUS-3 residues 41-259; p*I* of 5.5 for P22 residues 41-263.

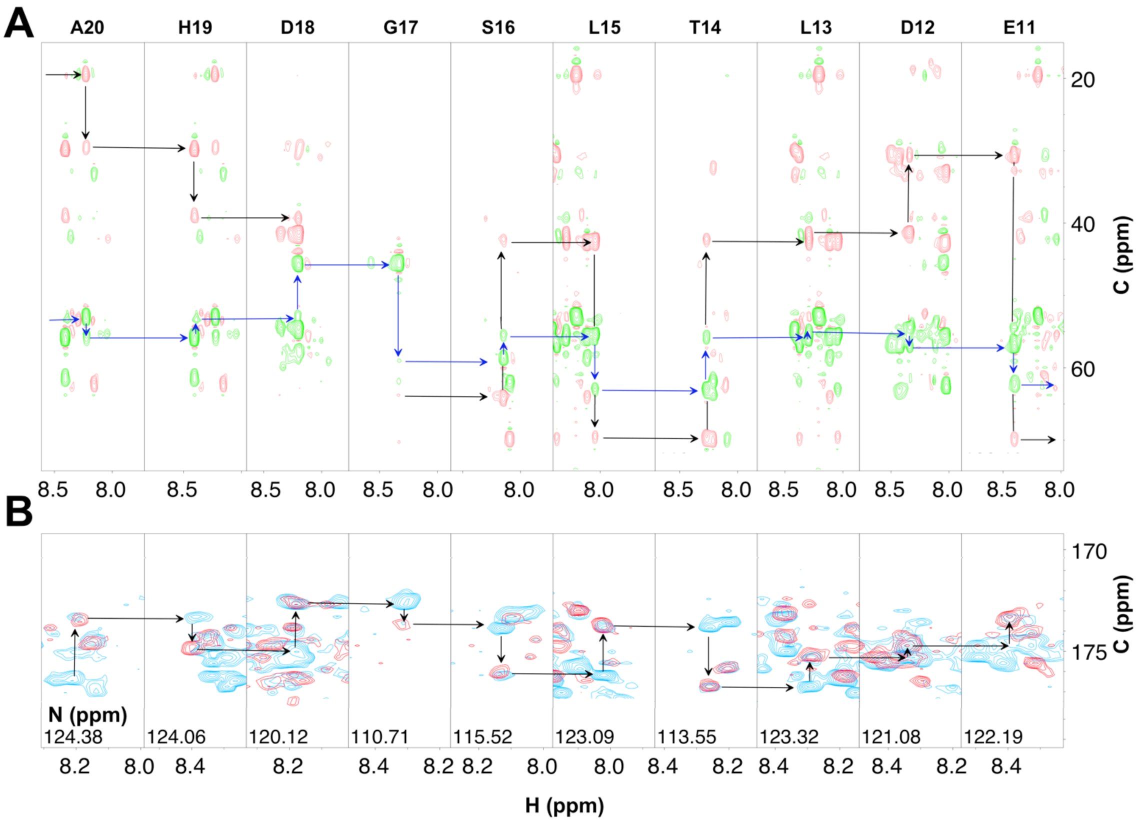

**Fig. S4 Representative strips from 3D NMR spectra illustrating sequential assignments of the A20-E11 segment of P22 His$_6$-SP. (A)** 3D HNCACB spectrum. Cα correlations are in green, Cβ correlations are in pink. The experiment establishes sequential connectivities through $^{13}$Cα (blue) and $^{13}$Cβ (black) nuclei. **(B)** Superposition of 3D HNCO (red) and HN(CA)CO (blue) spectra, illustrating connectivities through $^{13}$C' carbonyl atoms. NMR data were recorded on samples containing 0.5 mM $^{13}$C/$^{15}$N P22 His$_6$-SP, in 20 mM sodium phosphate (pH 6.0), at a temperature of 35 $^{o}$C.

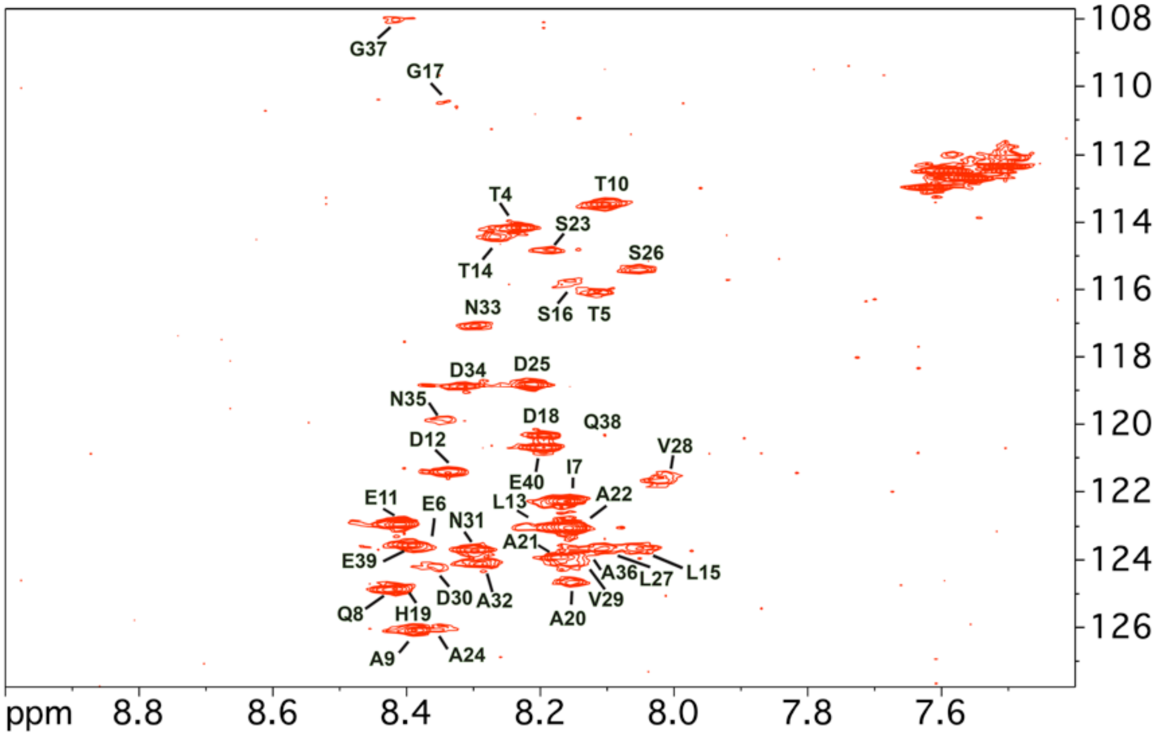

**Fig. S5 Assignments for $^{1}H$-$^{15}N$ HSQC correlations that persist when wild type P22 SP is encapsulated into capsids.** P22 SP (~300 µM) was encapsulated into empty phage P22 procapsids at a ratio of 60:1 to ensure tight binding (Parker et al., 2001), as described in the Methods. Samples contained 20 mM sodium phosphate (pH 7.0) and 50 mM NaCl to retain encapsulation conditions (Suhanovsky and Teschke, 2011). The temperature was 35 °C. The NMR data for capsid-encapsulated SP are the same as in Fig. 3B.

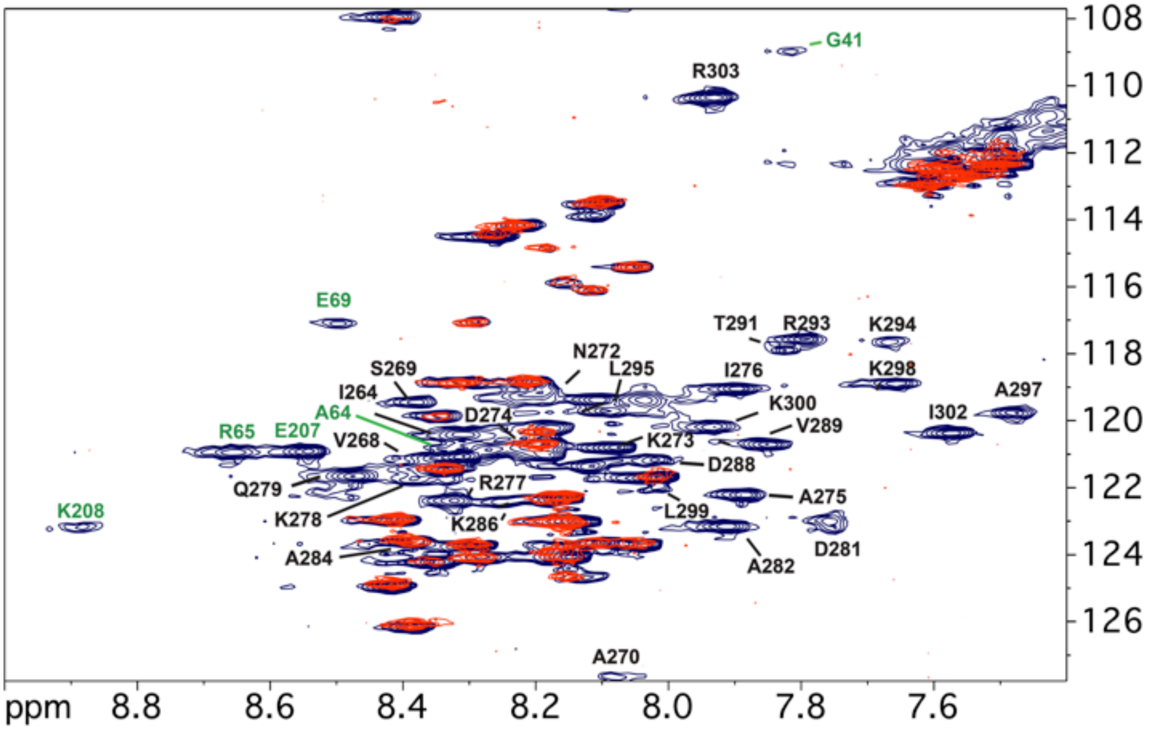

**Fig. S6 Assignments of $^{1}$H-$^{15}$N HSQC correlations from free wild type P22 SP.** The spectra shown to indicate assignments are the same as in Fig. 3B of the main text. NMR data for free 115 µM P22 SP in 20 mM sodium phosphate buffer (pH 7.0) were obtained at 35 $^{o}$C, under equivalent conditions to those in Fig. S5. Crosspeaks from the HTH domain are indicated with black labels, those from other regions of the protein with green labels. NMR assignments for the disordered N-terminal 4-40 segment seen in both free and encapsulated SP (red) are given in Fig. S5.

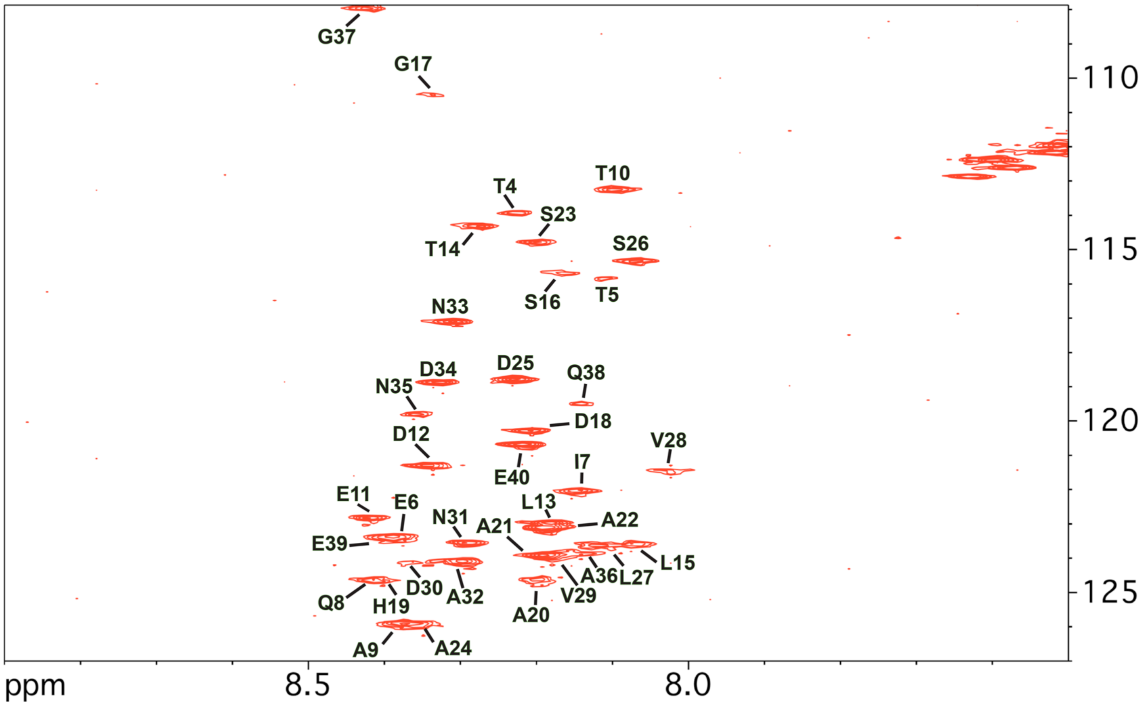

**Fig. S7 Assignments for $^{1}$H-$^{15}$N HSQC correlations that persist when P22 His$_6$-SP is encapsulated into capsids.** The data are analogous to those in Fig. S5, except using His$_6$-SP instead of the wild type SP without a His$_6$-tag. The similarity of this spectrum to that in Fig. S5, indicates that the affinity tag used for protein purification does not interfere with the experiments. P22 His$_6$-SP (~300 µM) was encapsulated into phage P22 procapsids at a ratio of 60:1. Samples contained 20 mM sodium phosphate (pH 7.0) and 50 mM NaCl. The temperature was 40 $^{o}$C.

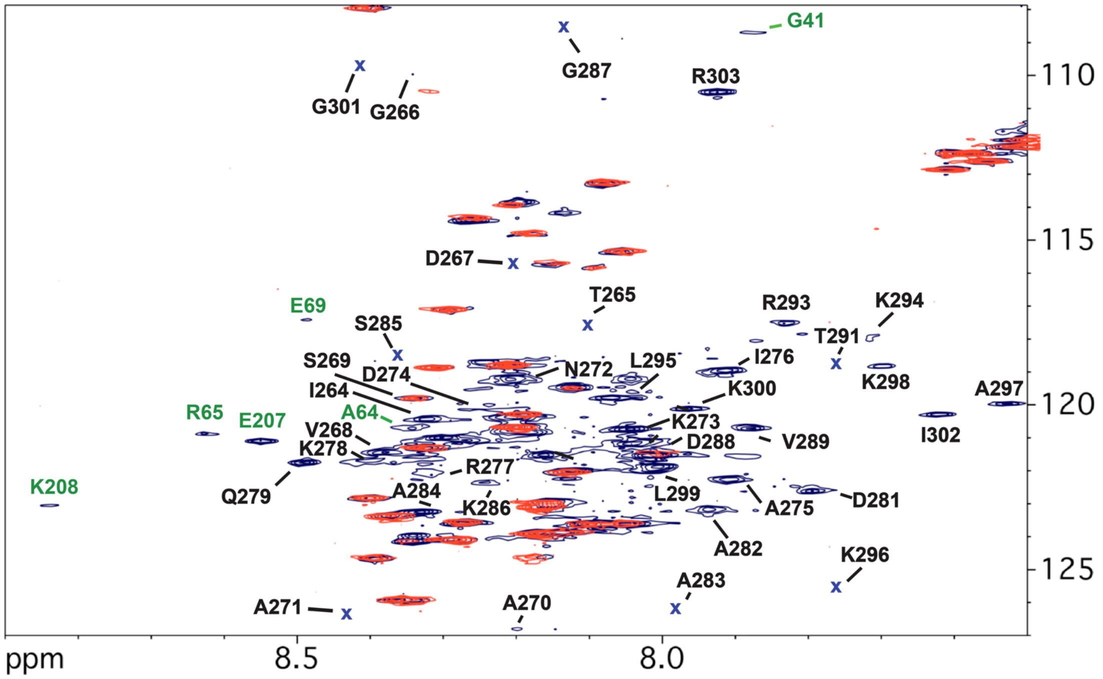

**Fig. S8 Assignments for $^{1}$H-$^{15}$N HSQC correlations from free P22 His$_6$-SP.** The data are analogous to those in Fig. S6 and Fig. 3B of the main text, except using His$_6$-SP instead of the wild type SP. The similarity of this spectrum to that in Fig. S6 indicates the affinity tag used for protein purification does not interfere with experiments. NMR spectra for free His$_6$-SP (blue) were obtained under equivalent conditions to the capsid-encapsulated His$_6$-SP (red) described in Fig. S7. Crosspeaks from the HTH domain are indicated with black labels, those from other regions of the protein with green labels. NMR assignments for the disordered N-terminal 4-40 segment seen in both free and encapsulated SP (red) are given in Fig. S7. Positions marked "x" correspond to crosspeaks seen in the 3D experiments used to obtain NMR assignments for His$_6$-SP, that were too weak to see at the displayed contour level in the 2D $^{1}$H-$^{15}$N HSQC spectrum.

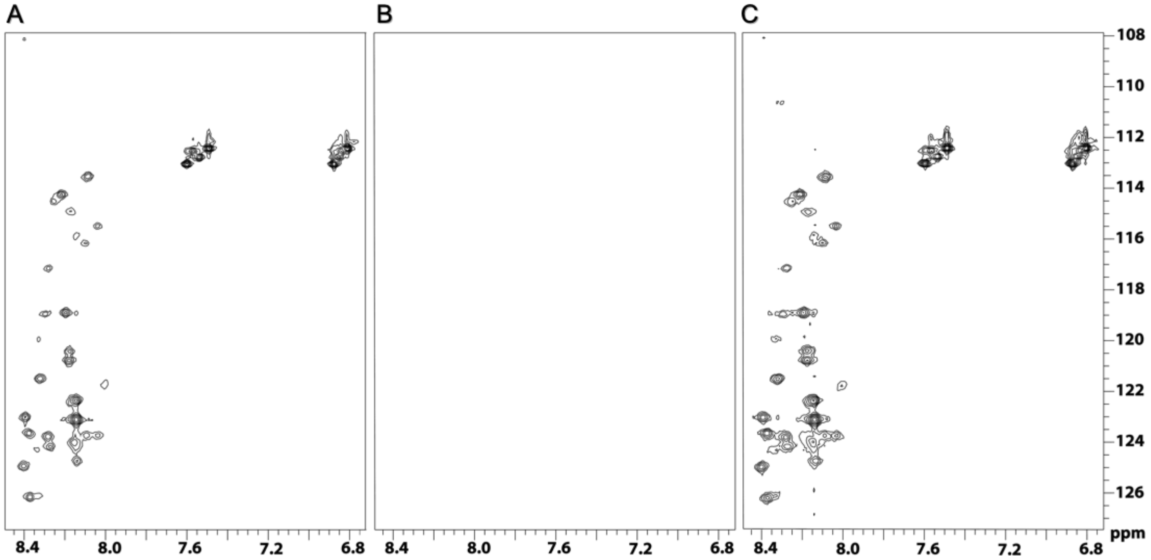

**Fig. S9 Control experiments to demonstrate encapsulation of wild type SP into capsids. (A)** SP encapsulated into P22 capsids, as in Fig. S5. **(B)** Spectrum of the supernatant obtained after sedimenting the SP-capsid complex at 104,813 g for 20 min. **(C)** Spectrum of the sedimented SP-capsid complex, resuspended in 20 mM sodium phosphate (pH 7.0) by shaking at 180 osc/min for 20 h at 4 $^{o}$C. Equivalent results were obtained when the control experiments were done with $His_6$-SP (not shown).

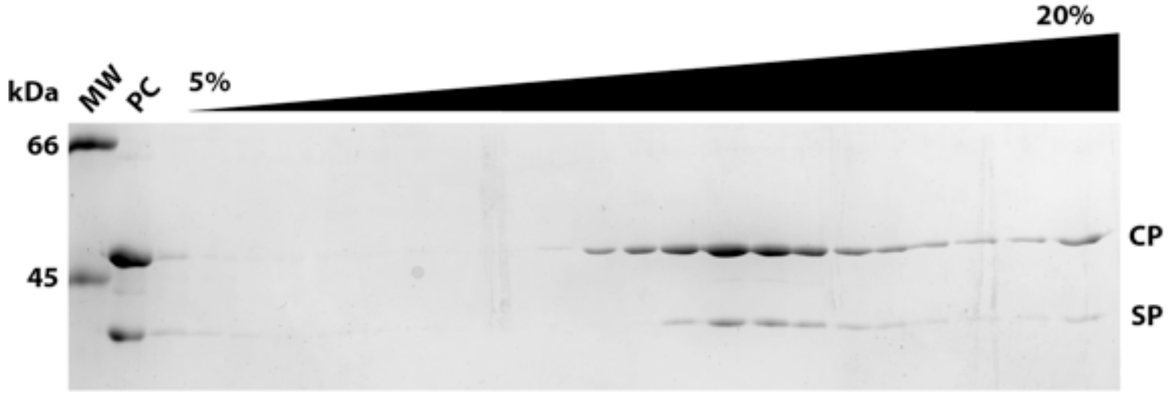

**Fig. S10 Control experiments verifying that wild type P22 SP is encapsulated in phage P22 procapsids.** P22 procapsids (PC) with encapsulated scaffolding protein (SP), prepared as described in Experimental Procedures, were run on a 5 to 20% sucrose gradient and fractions were visualized by 10% SDS-PAGE. The sucrose gradient was fractionated from the top and loaded left to right. Molecular weight marker is indicated by MW, and a procapsid marker is indicated by PC. The constituent coat protein (CP) and scaffolding protein (SP) of the PC marker are indicated on the right side of the gel. That CP and SP co-sediment, and the absence of free SP (empty lanes in the beginning of the gradient) demonstrates that SP is encapsulated in the PCs.

**Table S1: NMR Assignments for the P22 SP.**

| | Chemical Shift (ppm)[a] | | | | | | |
|---|---|---|---|---|---|---|---|
| Residue | N | HN | C | Cα | Cβ | Hα | Others |
| Pro 3 | | - | 175.91 | 62.88 | 33.74 | | |
| Thr 4 | 113.63 | 8.20 | 173.58 | 61.89 | 69.56 | 4.50 | Hβ 4.31; Hγ2Me 1.31 |
| Thr 5 | 115.58 | 8.10 | 173.14 | 61.89 | 69.46 | 4.43 | Hγ2Me 1.23 |
| Glu 6 | 123.14 | 8.38 | 175.01 | 56.73 | 30.73 | | |
| Ile 7 | 121.76 | 8.13 | 174.71 | 61.38 | 38.63 | | |
| Gln 8 | 124.44 | 8.40 | 174.15 | 56.10 | 29.9 | | |
| Ala 9 | 125.68 | 8.37 | 176.48 | 52.54 | 19.68 | | |
| Thr 10 | 113.06 | 8.09 | 173.31 | 61.89 | 69.81 | 4.35 | Hγ 1.26 |
| Glu 11 | 122.60 | 8.41 | 174.70 | 57.11 | 30.81 | | |
| Asp 12 | 121.08 | 8.33 | 175.27 | 54.35 | 41.19 | 4.72 | Hβ 2.66 |
| Leu 13 | 122.69 | 8.16 | 176.77 | 55.79 | 43.87 | 4.17 | Hβ 1.85; HδMe 0.91 |
| Thr 14 | 114.17 | 8.27 | 173.88 | 62.82 | 69.38 | 4.27 | Hβ 4.03; Hγ2Me 1.27 |
| Leu 15 | 123.40 | 8.06 | 176.22 | 55.33 | 42.09 | | |
| Ser 16 | 115.52 | 8.16 | 173.66 | 59.00 | 63.77 | 4.50 | Hβ 3.95 |
| Gly 17 | 110.40 | 8.32 | 172.42 | 45.83 | - | | Hα 4.03 |
| Asp 18 | 120.12 | 8.21 | 174.82 | 53.33 | 39.27 | | |
| His 19 | 124.37 | 8.40 | 173.53 | 55.33 | 29.57 | | |
| Ala 20 | 124.38 | 8.17 | 176.46 | 52.53 | 19.70 | | |
| Ala 21 | 123.66 | 8.18 | 176.41 | 53.42 | 19.63 | | |
| Ala 22 | 122.81 | 8.16 | 176.47 | 52.42 | 19.55 | 4.46 | HβMe 1.45 |
| Ser 23 | 114.58 | 8.18 | 173.44 | 58.51 | 63.77 | 4.49 | Hβ 3.94 |
| Ala 24 | 125.65 | 8.36 | 176.40 | 52.53 | 19.49 | 4.33 | |
| Asp 25 | 118.60 | 8.21 | 176.11 | 54.72 | 41.45 | 4.71 | Hβ 2.69 |
| Ser 26 | 115.14 | 8.06 | 173.00 | 58.81 | 63.84 | 4.49 | Hβ 3.91 |
| Leu 27 | 123.38 | 8.11 | 175.63 | 55.35 | 42.58 | 4.40 | Hβ 1.67; HδMe 0.95 |
| Val 28 | 121.32 | 8.02 | 174.76 | 62.67 | 32.81 | 4.14 | Hβ 2.09; HγMe 0.96 |
| Val 29 | 123.72 | 8.15 | 174.30 | 62.18 | 33.06 | 4.20 | Hβ 2.12; HγMe 0.97 |
| Asp 30 | 123.90 | 8.35 | 175.23 | 54.23 | 41.50 | 4.74 | |
| Asn 31 | 123.32 | 8.29 | 175.32 | 55.59 | 42.16 | | |
| Ala 32 | 123.86 | 8.29 | 176.52 | 53.60 | 19.62 | 4.29 | Hβ 1.43 |
| Asn 33 | 116.89 | 8.30 | 173.84 | 53.47 | 39.31 | 4.75 | Hβ 2.83 |
| Asp 34 | 118.63 | 8.32 | 173.88 | 53.41 | 39.35 | 4.75 | Hβ 2.86 |
| Asn 35 | 119.57 | 8.35 | 174.73 | 53.40 | 39.32 | 4.75 | Hβ 2.83 |
| Ala 36 | 123.64 | 8.15 | 177.02 | 53.43 | 19.55 | 4.32 | HβMe 1.47 |
| Gly 37 | 107.73 | 8.42 | 173.02 | 45.87 | - | | Hα 3.99 |
| Gln 38 | 119.23 | 8.13 | 174.78 | 56.12 | 29.96 | 4.40 | Hβ 2.03; Hγ 2.36 |

| | | | | | | | |
|---|---|---|---|---|---|---|---|
| Glu 39 | 123.14 | 8.39 | 173.83 | 54.30 | 30.79 | 4.32 | Hβ 2.00; Hγ 2.28 |
| Glu 40 | 120.46 | 8.21 | 173.71 | 54.49 | 28.81 | 4.25 | Hβ 1.98; Hγ 2.23 |
| *Gly 41* [b] | 108.57 | 7.84 | 170.34 | 45.93 | - | | |
| *Phe 63* [b] | - | - | 176.48 | 57.31 | 38.35 | | |
| *Ala 64* [b] | 120.78 | 8.31 | 175.90 | 54.09 | 18.96 | | |
| *Arg 65* [b] | 120.64 | 8.64 | - | 58.16 | 30.01 | | |
| *Ile 68* [b] | - | - | 174.69 | 63.75 | 41.25 | | |
| *Glu 69* [b] | 117.24 | 8.47 | - | 59.96 | 30.72 | | |
| *Pro 206* [c] | - | - | 176.42 | 65.41 | 32.39 | | |
| *Glu 207* [c] | 120.80 | 8.56 | - | 59.77 | 29.95 | 4.67 | Hβ 2.7 |
| *Lys 208* [c] | 122.98 | 8.89 | 176.20 | 59.35 | 33.70 | 5.76 | |
| Pro 263 | - | - | 174.77 | 63.36 | 32.72 | | |
| Ile 264 | 120.55 | 8.32 | 175.30 | 61.37 | 38.72 | | |
| Thr 265 | 117.27 | 8.10 | 173.65 | 62.04 | 70.07 | 3.85 | |
| Gly 266 | 110.58 | 8.34 | 172.66 | 45.64 | - | | Hα 4.01 |
| Asp 267 | 115.81 | 8.24 | 174.98 | 54.38 | 41.30 | | |
| Val 268 | 120.44 | 8.34 | 174.90 | 61.01 | 32.82 | | |
| Ser 269 | 119.56 | 8.38 | 172.26 | 58.18 | 63.94 | | |
| Ala 270 | 126.55 | 8.22 | 173.65 | 50.60 | 18.63 | 4.16 | HβMe 1.47 |
| Ala 271 | 126.93 | 8.37 | 176.16 | 52.66 | 19.53 | 4.46 | HβMe 1.44 |
| Asn 272 | 119.08 | 8.20 | 174.58 | 54.00 | 41.24 | 4.70 | Hβ 2.42 |
| Lys 273 | 120.59 | 8.05 | 175.57 | 56.82 | 30.16 | | Hβ 1.97 |
| Asp 274 | 120.20 | 8.31 | 173.29 | 55.31 | 41.11 | 4.71 | Hβ 2.74 |
| Ala 275 | 121.94 | 7.91 | 177.89 | 54.22 | 18.76 | 3.89 | HβMe 1.40 |
| Ile 276 | 118.84 | 7.91 | 175.91 | 64.69 | 37.51 | 4.10 | Hb 1.90; Hγ1 1.52; Hγ2Me 0.95 |
| Arg 277 | 121.48 | 8.32 | 175.34 | - | 30.78 | | |
| Lys 278 | 121.64 | 8.43 | 174.75 | 56.87 | 30.86 | | |
| Gln 279 | 121.53 | 8.49 | 175.28 | 56.20 | 30.05 | 4.32 | Hβ 2.02; Hγ 2.28 |
| Met 280 | | | 174.47 | 55.24 | 32.82 | | |
| Asp 281 | 122.59 | 7.77 | 173.35 | 52.83 | 42.41 | | |
| Ala 282 | 122.97 | 7.93 | 175.79 | 54.35 | 18.78 | 4.49 | HβMe 1.66 |
| Ala 283 | 126.19 | 7.98 | | 50.80 | 18.67 | | |
| Ala 284 | 123.01 | 8.33 | 176.20 | 54.35 | 19.64 | 4.40 | HβMe 1.45 |
| Ser 285 | 118.43 | 8.40 | 173.72 | 59.17 | 63.87 | | |
| Lys 286 | 122.62 | 8.40 | | 57.10 | 32.22 | | |
| Gly 287 | 107.09 | 7.94 | 172.44 | 45.95 | - | | Hα 2.65 |

| | | | | | | | |
|---|---|---|---|---|---|---|---|
| Asp 288 | 120.95 | 8.04 | 173.07 | 55.26 | 42.70 | 4.17 | Hβ 3.06 |
| Val 289 | 120.50 | 7.87 | 177.16 | 59.78 | 32.78 | | |
| Glu 290 | | | | 54.22 | 34.55 | | |
| Thr 291 | 118.54 | 7.75 | 172.44 | 59.01 | 70.25 | | |
| Tyr 292 | | | 177.91 | 59.58 | 41.29 | | |
| Arg 293 | 117.43 | 7.83 | 176.75 | - | 30.27 | 4.41 | Hβ 1.88 |
| Lys 294 | 117.66 | 7.69 | 175.87 | 57.28 | 32.91 | | |
| Leu 295 | 119.54 | 8.08 | - | 55.28 | 42.14 | 4.14 | Hβ 1.97; HγMe 0.71 |
| Lys 296 | 125.74 | 7.72 | 174.69 | 58.13 | 34.15 | 4.19 | Hβ 1.79; Hγ 1.41 |
| Ala 297 | 119.69 | 7.50 | 175.34 | 54.62 | 18.62 | 4.16 | HβMe 1.19 |
| Lys 298 | 118.66 | 7.67 | - | 58.39 | 32.79 | | |
| Leu 299 | 121.94 | 8.04 | 174.31 | 55.52 | 43.12 | | |
| Lys 300 | 120.19 | 8.00 | 174.75 | 56.22 | 30.89 | | |
| Gly 301 | 110.12 | 8.44 | 172.49 | 45.77 | - | | Hα 3.96 |
| Ile 302 | 120.15 | 7.60 | 173.92 | 61.61 | 38.57 | 3.59 | Hβ 1.40 |
| Arg 303 | 110.15 | 7.94 | 170.43 | 57.89 | 31.75 | 4.19 | Hβ 1.87 |

[a] Chemical shift assignments were obtained from 3D NMR experiments (described in Methods) recorded on 0.5 mM $^{13}C/^{15}N$-labeled samples of P22 $His_6$-SP in 20 mM sodium phosphate buffer at pH 6.0, and a temperature of 35 $^oC$.

[b] Tentative assignment due to isolated spin systems in the amino acid sequence

[c] The tentative sequence-specific assignments for the segment P206-E207-K208, could alternatively belong to the segment P220-E221-K222.

### Supporting Materials Reference

Casjens, S., Winn-Stapley, D.A., Gilcrease, E.B., Morona, R., K√ºhlewein, C., Chua, J.E.H., Manning, P.A., Inwood, W., and Clark, A.J. (2004). The Chromosome of Shigella flexneri Bacteriophage Sf6: Complete Nucleotide Sequence, Genetic Mosaicism, and DNA Packaging. Journal of Molecular Biology *339*, 379-394.

Casjens, S.R., and Thuman-Commike, P.A. (2011). Evolution of mosaically related tailed bacteriophage genomes seen through the lens of phage P22 virion assembly. Virology *411*, 393-415.

Davison, A.J. (2011). Evolution of sexually transmitted and sexually transmissible human herpesviruses. Annals of the New York Academy of Sciences *1230*, E37-49.

Dokland, T. (1999). Scaffolding proteins and their role in viral assembly. Cellular & Molecular Life Sciences *56*, 580-603.

King, M.R., Vimr, R.P., Steenbergen, S.M., Spanjaard, L., Plunkett, G., 3rd, Blattner, F.R., and Vimr, E.R. (2007). Escherichia coli K1-specific bacteriophage CUS-3 distribution and function in phase-variable capsular polysialic acid O acetylation. J Bacteriol *189*, 6447-6456.

Parker, M.H., Brouillette, C.G., and Prevelige, P.J. (2001). Kinetic and calorimetric evidence for two distinct scaffolding protein binding populations within the bacteriophage P22 procapsid. Biochemistry *40*, 8962-8970.

Pedulla, M.L., Ford, M.E., Karthikeyan, T., Houtz, b.M., Hendrix, R.W., Hatfull, G.F., Poteete, A.R., Gilcrease, E.B., Winn-Stapley, D.A., and Casjens, S.R. (2003). Corrected sequence of the bacteriophage p22 genome. J. Bacteriol. *185*, 1475-1477.

Suhanovsky, M.M., and Teschke, C.M. (2011). Bacteriophage P22 capsid size determination: Roles for the coat protein telokin-like domain and the scaffolding protein amino-terminus. Virology *417*, 418-429.
